# The histone variant H2A.W and linker histone H1 co-regulate heterochromatin accessibility and DNA methylation

**DOI:** 10.1101/2020.03.19.998609

**Authors:** Pierre Bourguet, Colette L. Picard, Ramesh Yelagandula, Thierry Pélissier, Zdravko J. Lorković, Suhua Feng, Marie-Noëlle Pouch-Pélissier, Anna Schmücker, Steven E. Jacobsen, Frédéric Berger, Olivier Mathieu

## Abstract

In flowering plants, heterochromatin is demarcated by the histone variant H2A.W, elevated levels of the linker histone H1, and specific epigenetic modifications, such as high levels of DNA methylation at both CG and non-CG sites. How H2A.W regulates heterochromatin organization and interacts with other heterochromatic features is unclear. To analyze the *in vivo* function of H2A.W, we created a *h2a.w* null mutant via CRISPR-Cas9, *h2a.w-2.* We find that H2A.W antagonizes deposition of H1 at heterochromatin and that non-CG methylation and accessibility are moderately decreased in *h2a.w-2* heterochromatin. Compared to H1 loss alone, combined loss of H1 and H2A.W greatly increases accessibility and facilitates non-CG DNA methylation in heterochromatin, suggesting co-regulation of heterochromatic features by H2A.W and H1. Our results suggest that H2A.W helps maintain optimal heterochromatin accessibility and DNA methylation by promoting chromatin compaction together with H1, while also inhibiting excessive H1 incorporation.

## Introduction

Eukaryotic genomes are packaged in chromatin. The basic unit of chromatin is the nucleosome, which contains a protein octamer comprising two of each of the core histones H2A, H2B, H3, and H4, wrapped by ~147bp of DNA. Chromatin is organized into two distinct domains termed constitutive heterochromatin, which is enriched in transposable elements (TEs) and other types of repetitive DNA, and euchromatin, which comprises mostly protein-coding genes. Euchromatin is more accessible and associated with transcriptional activity, whereas heterochromatic domains prevent transcription and are often compacted into higher order structures such as chromocenters in *Arabidopsis thaliana* and mouse nuclei. Yet, heterochromatin has to retain a certain degree of accessibility to allow important DNA-related biological processes to occur, including maintenance of DNA methylation, DNA replication, DNA damage repair, and transcription for small RNA production.

In most eukaryotes, euchromatic and heterochromatic regions can be distinguished by their DNA methylation level, the presence of distinct post-translational modifications of histones, and their association with specific histone variants. In plants, DNA methylation occurs in three sequence contexts (CG, CHG, and CHH, where H is any base but G). DNA methylation in euchromatic regions tends to be low, except at CG sites over gene bodies of protein-coding genes. Heterochromatic sequences, however, are characterized by dense methylation at all three sequence contexts (CG and non-CG methylation). In *Arabidopsis,* heterochromatin is additionally decorated with histone H3 lysine 9 mono and dimethylation (H3K9me1 and H3K9me2), catalyzed by the SU(VAR)3–9 HOMOLOG-class of histone methyltransferases SUVH4/KYP, SUVH5, and SUVH6^1^. The mechanisms that establish and maintain heterochromatin-specific non-CG methylation and H3K9 methylation are tightly and reciprocally interconnected^2–5^. The DNA methyltransferases CMT2 and CMT3 are recruited by H3K9me1 and H3K9me2^2^, while the H3K9 methyltransferases SUVH4/5/6 are recruited to chromatin by DNA methylation^6–8^, creating a positive feedback loop that reinforces silencing. DNA methylation in plants is also established and maintained via the RNA-directed DNA methylation (RdDM) pathway, which preferentially targets short euchromatic TEs and the edges of long heterochromatic TEs^2,3,9^ Recruitment of RNA polymerase IV during the early steps of RdDM involves SHH1, which binds methylated H3K9, thereby also linking RdDM targeting to H3K9 methylation^10,11^. *Arabidopsis* heterochromatin is also marked by H3K27me1, which depends on the redundant histone methyltransferases ARABIDOPSIS TRITHORAX-RELATED 5 and 6 (ATXR5, ATXR6)^12^. Although heterochromatin structure is visibly altered in *atxr5 atxr6* mutants, DNA methylation and H3K9me2 appear largely unaffected, suggesting that H3K27me1 is maintained independently of these marks.

The linker histone H1, which binds nucleosomes and the intervening linker DNA, is also preferentially associated with heterochromatin in *Arabidopsis*^13–17^. In *Arabidopsis,* H1 associates with chromatin independently of DNA methylation, but loss of H1 leads to chromocenter decondensation, and has varying effects on DNA methylation: pericentromeric heterochromatic TEs gain DNA methylation in *h1*, while TEs on the chromosome arms lose methylation^15,18–20^. H1 is thought to hinder heterochromatic DNA methylation by restricting the access of DNA methyltransferases to these regions^3,2^. The joint action of H1 and CG methylation by the DNA methyltransferase MET1 silences a subset of TEs and prevents the production of aberrant gene transcripts, suggesting DNA methylation and H1 help define functional transcriptional units^15^. The histone variant H3.3 also plays a role in restricting H1 from associating with active genes in *Arabidopsis*^17^, Drosophila^22^, and mouse^23^. However, the mechanisms that control H1 deposition and shape its relative enrichment in heterochromatin remain unknown.

In *Arabidopsis,* the histone variant H2A.W is strictly and specifically localized to constitutive heterochromatin^24,25^. H2A variants comprise the most diverse histone family and directly impact biochemical properties of the nucleosome^26,27^. In *Arabidopsis,* nucleosomes typically contain a single type of H2A variant – either replicative H2A, H2A.X, H2A.Z, or H2A.W^28^. In land plants, the majority of H2A.Z nucleosomes associate with repressive H3K27me3 modifications while a small fraction associates with active H3K36me3 marks found at the transcription start site in transcribed genes^24,29^. Most of the body of expressed genes is occupied by replicative H2A and H2A.X nucleosomes^24^. The exclusive localization of H2A.W at constitutive heterochromatin is unique among Arabidopsis H2A variants^24,30^. While specialized histone chaperones mediate the incorporation of specific H2A variants, no chaperone dedicated to H2A.W has been identified so far^27,31^. *Arabidopsis* has three H2A.W isoforms, H2A.W.6, H2A.W.7, and H2A.W.12, encoded by *HTA6, HTA7,* and *HTA12* respectively. To characterize the role of H2A.W in *Arabidopsis* heterochromatin, a previous study generated a triple-knockout *hta6 hta7 hta12* line, referred here to as *h2a.w-1*^24^

However, we identified a large genomic rearrangement in the *hta6* transfer-DNA (T-DNA) insertion mutant allele used to generate the *h2a.w-1* triple knockout line. Using CRISPR-Cas9, we obtained a new null *h2a.w* triple mutant without this rearrangement, referred to here as *h2a.w-2*. Analyzing *h2a.w-2* mutants revealed that the *hta6* chromosomal rearrangement, which led to a duplication of the *CMT3* locus, was responsible for the severe developmental effects and CHG hypermethylation reported in *h2a.w-1*^24^. These defects hindered functional analysis of H2A.W. Using the new mutant *h2a.w-2,* we now show that loss of H2A.W results in no visible developmental or morphological phenotypes and has only minor effects on gene and TE expression. In *h2a.w-2,* heterochromatin exhibits a mild decrease in accessibility accompanied by increased deposition of replicative H2A and H2A.X and of the linker histone H1, as well as decreased levels of non-CG methylation. The combined loss of H1 and H2A.W enhances both chromatin accessibility and DNA methylation in pericentromeric heterochromatin. Based on these results, we propose that H2A.W and H1 jointly regulate DNA methylation and heterochromatin accessibility.

## Results

### A large chromosomal translocation in *hta6* obscured functional analysis of H2A.W

To investigate whether H2A.W plays a role in controlling TE mobilization, we used available Whole Genome Bisulfite Sequencing (BS-seq) data of *h2a.w-1* triple mutants and their corresponding wild type (WT)^24^. We detected significantly increased copy numbers for several TEs (Supplementary Table 1), but because these TEs were all located within the same genomic region on the right arm of chromosome 1, we suspected that this result reflected a chromosomal rearrangement in the *h2a.w-1* plants, rather than a genuine role for H2A.W in controlling activity of this TE subset. Further analysis of *h2a.w-1* BS-seq and RNA-seq data revealed abnormally high coverage along an approximately 5 Mb region of chromosome 1, indicating that this region may be duplicated in *h2a.w-1* (Fig. 1a, Supplementary Fig. 1a). Southern blot analysis of *hta6* SALK_024544.32 (hereafter named *hta6-1), hta7* (GABI_149G05.01), and *hta12* (SAIL_667_D09) DNA confirmed the presence of a genomic rearrangement, likely a translocation of part of chromosome 1, in *hta6-1* (Fig. 1b). Further analyses revealed a ~5 Mb deletion in chromosome 1 which is replaced by T-DNA/vector sequences, and a translocation of a part of chromosome 1, flanked by T-DNA sequences, to chromosome 5 (Supplementary Fig. 1b). Using segregating plants from crosses between wild-type and *hta6-1,* we were able to recover *hta6-1* mutants with either normal or doubled dosage of the chromosome 1 region. The plants with normal dosage showed a WT-like phenotype, while plants with doubled dosage were abnormally small (Supplementary Fig. 1b), suggesting that increased dosage of this portion of chromosome 1 causes developmental defects.

**Figure 1.**
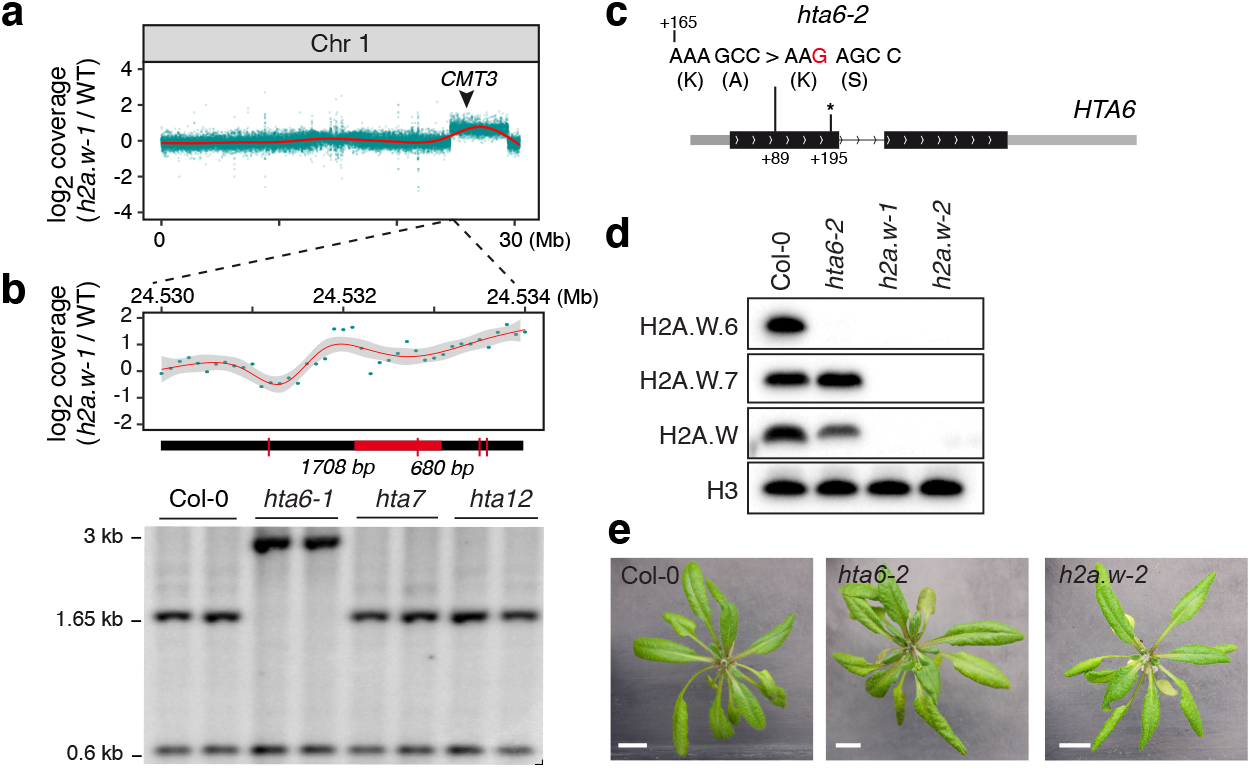
H2A.W is not required for *Arabidopsis* development. (a) Sequencing coverage of published *h2a.w-1* BS-seq data^24^, averaged in 1 kb bins across chromosome 1. The red line shows the smoothed conditional mean (LOESS). The black arrowhead indicates the genomic location of *CMT3.* See also Supplementary Fig 1a. (b) Zoomed-in view of plot in (a) across the left border of the chromosome 1 region showing increased coverage in *h2a.w-1* (top panel). DNA gel blot analysis of the chromosome 1 region showing abnormal coverage in *h2a.w-1* in the indicated T-DNA insertion mutants (lower panel). Genomic DNA of the indicated genotypes was digested with *Ssp*I (recognition sites indicated by red ticks on the thick black line) and hybridized with a fragment corresponding to the genomic region indicated in red under the plot. (c) The *hta6-2* CRISPR-Cas9 mutant allele. Diagram of the *HTA6* gene showing the insertion of a G (in red) in *hta6-2*, which creates a frame shift 89 bp downstream from the translation initiation site and an early stop codon (asterisk) 195 bp downstream from the translation initiation site. (d) Western blot showing total loss of H2A.W in *h2a.w-1* and *h2a.w-2.* Nuclear extracts of the indicated genotypes were analyzed using antibodies directed against H2A.W.6, H2A.W.7, total H2A.W, and H3. (e) Representative images of wild-type, *hta6-2,* and *h2a.w-2* plants (scale bar = 1 cm). Both *hta6-2* and *h2a.w-2* mutants develop like wild-type Col-0 plants.

Using targeted mutagenesis via CRISPR-Cas9, we generated a new *hta6* allele *(hta6-2)* carrying a single-base frame shift mutation that causes a stop codon early in the protein (Fig. 1c). Western blot confirmed that *hta6-2* is a null mutant for H2A.W.6 (Fig. 1d). We crossed *hta6-2* with *hta7* and *hta12* to obtain the new null triple mutant, *h2a.w-2.* Western blot analysis confirmed that H2A.W is completely absent in *h2a.w-2* (Fig. 1d and Supplementary Fig. 1c). *h2a.w-2* plants are morphologically indistinguishable from wild-type plants (Fig. 1e), indicating that increased dosage of a large portion of chromosome 1, and not loss of H2A.W, caused the strong developmental defects reported in *h2a.w-1*^24^. Instead, H2A.W appeared to be dispensable for *Arabidopsis* development. We therefore sought to clarify the function of H2A.W in heterochromatin transcription, composition, and organization.

### H2A.W has little impact on transcription but is required for the efficient methylation of heterochromatic DNA

Previous analyses of the impact of H2A.W loss on transcription may have been confounded by the genomic rearrangement in *h2a.w-1.* We therefore re-explored whether lack of H2A.W affects genome-wide transcription by performing RNA-seq of WT and *h2a.w-2.* These analyses identified only a few differentially expressed protein-coding genes (PCGs; 78 upregulated, 52 downregulated) and only a handful of transcriptionally activated TEs in *h2a.w-2* (Supplementary Fig. 2a, b). These results show that gene expression is not strongly affected in the absence of H2A.W, and that H2A.W either does not play a significant role in repressing TEs or that other pathways are compensating for H2A.W loss.

The chromosomal rearrangement in *h2a.w-1* caused a duplication of the gene encoding CMT3 (Fig. 1a, Supplementary Fig. 1a) and this duplication was likely responsible for the higher levels of CHG methylation previously reported in *h2a.w-1*^24^. Indeed, we found that *CMT3* mRNA was significantly increased in *hta6-1* and in *h2a.w-1* carrying additional *CMT3* gene copies (Supplementary Fig. 2c). However, *CMT3* expression in *h2a.w-2* was similar to WT (Supplementary Fig. 2c), indicating that *CMT3* transcription is not affected by loss of H2A.W. We were also able to recover *h2a.w-2 cmt3* quadruple mutant plants, which had no obvious developmental defects (Supplementary Fig. 2d), indicating that the lethal genetic interaction between *h2a.w-1* and *cmt3*^24^ was due to the chromosomal rearrangement in *hta6-1.*

We therefore sought to clarify the impact of H2A.W loss on DNA methylation by examining the methylome of *h2a.w-2* using BS-seq. We found no conspicuous change in CG DNA methylation in the *h2a.w-2* mutant (Supplementary Fig. 3a, b), while non-CG methylation levels appeared substantially decreased at pericentromeric regions (Supplementary Fig. 3a). In agreement with these chromosome-wide observations, TEs located in the pericentromeres showed partially reduced CHG and CHH methylation levels (Fig. 2a, Supplementary Fig. 3c), suggesting that H2A.W is required for maintenance of DNA methylation in these regions. Conversely, we found that TEs located on chromosome arms showed increased CHH DNA methylation in *h2a.w-2,* suggesting an antagonistic effect of H2A.W (Fig. 2a, Supplementary Fig. 3c). Indeed, looking specifically at regions normally occupied by H2A.W in WT revealed opposing changes in non-CG DNA methylation in the *h2a.w-2* mutant, based on chromosomal location. A substantial decrease in non-CG methylation levels was observed over H2A.W peaks in pericentromeric regions, whereas regions located in chromosome arms showed increased methylation levels (Fig. 2b, Supplementary Fig. 3d). Short TEs enriched in chromosome arms are known targets of the RdDM pathway involving DRM1/2, while CHG and CHH methylation at long heterochromatic TEs is preferentially maintained by CMT3 and CMT2, respectively^2,3^. We found that short TEs tended to gain CHH methylation in *h2a.w-2*, while long heterochromatic TEs instead tended to lose CHH and CHG methylation (Fig. 2c). Accordingly, non-CG methylation was reduced at CMT2-dependent regions but increased at DRM1/2-dependent regions in *h2a.w-2* (Fig. 2d). Together, these findings indicate that H2A.W promotes CMT3 and/or CMT2 mediated methylation maintenance in pericentromeric heterochromatin. However, H2A.W incorporation does not require CHG methylation^24^. Our data also suggest that H2A.W opposes RdDM at less heterochromatic regions on chromosome arms, supporting the idea that RdDM is inhibited by heterochromatin as proposed previously^3,32^.

**Figure 2.**
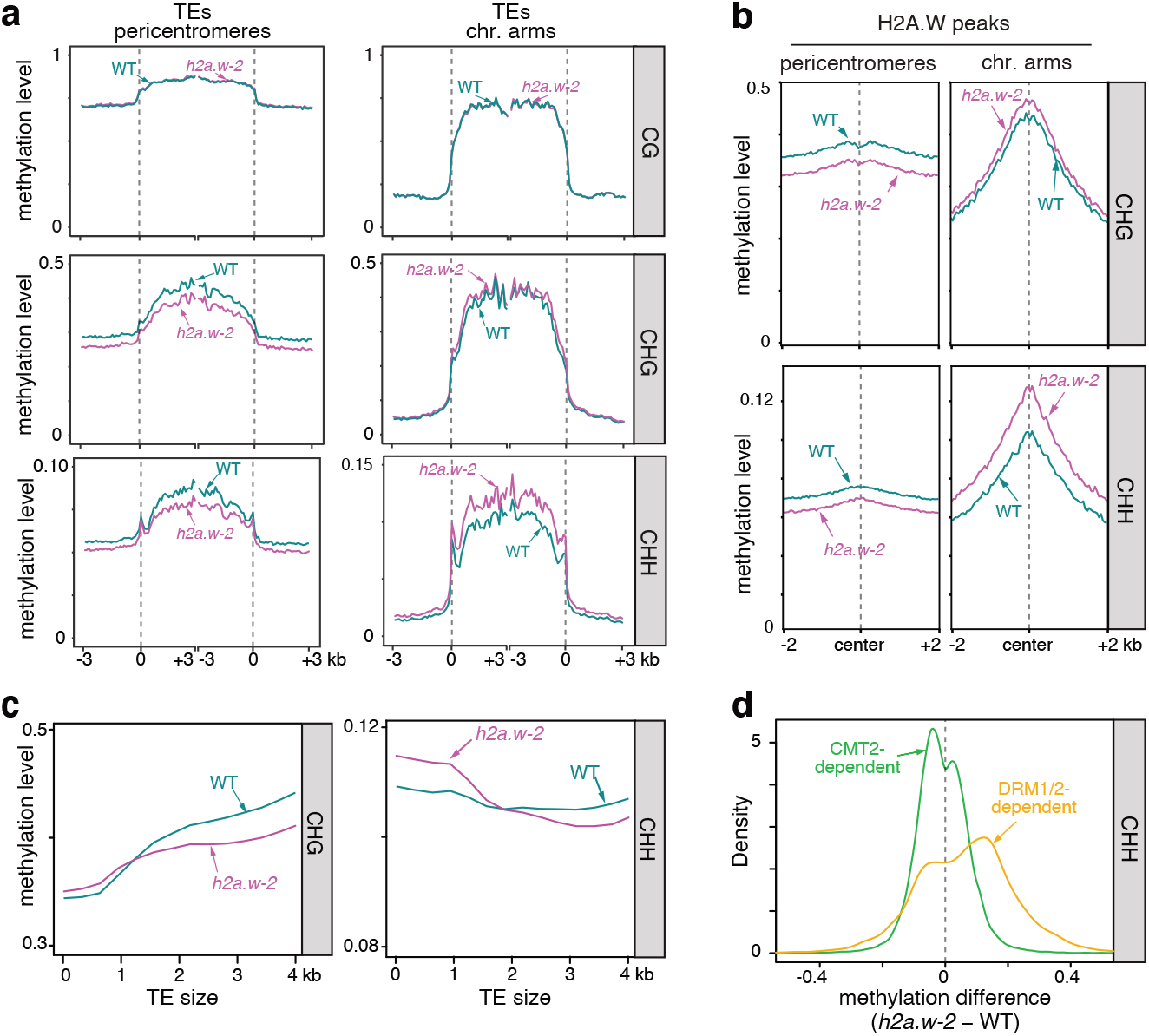
Loss of H2A.W alters non-CG DNA methylation patterns. (a) CG, CHG, and CHH methylation levels over TEs in pericentromeres and chromosome arms in WT and *h2a.w-2.* TEs were aligned at the 5’ (left dashed line) and 3’ end (right dashed line), and sequences 3 kb upstream or downstream were included, respectively. Average methylation over 100 bp bins is plotted. (b) CHG and CHH methylation levels over H2A.W peaks in the chromosome arms and pericentromeres. (c) Locally weighted scatterplot smoothing (LOESS) fit of CHG and CHH methylation levels in WT and *h2a.w-2* calculated in 50 bp windows and plotted against TE size. (d) Kernel density plots of CHH DNA methylation differences between *h2a.w-2* and WT at CMT2-dependent and DRM1/2-dependent regions.

### Heterochromatin accessibility decreases in the absence of H2A.W

Since complex and global changes in patterns of DNA methylation are often associated with modulation of chromatin accessibility^21^, we analyzed chromatin organization in *h2a.w-2.* Indeed, we observed enlarged chromocenters in *h2a.w-2* (Fig. 3a), as was previously observed in *h2a.w-1*^24^ Chromocenter enlargement is also observed in mutants that induce over-replication of heterochromatic regions^33^. To test for over-replication, we analyzed DNA content in *h2a.w-2* nuclei by FACS. We did not observe any obvious change relative to WT (Supplementary Fig. 4), indicating that enlargement of chromocenters most likely reflected changes in chromatin organization caused by loss of H2A.W, as reported previously^24^ By introducing a H2A.W.6 genomic construct in *h2a.w-2* mutants, we could partially rescue the enlargement of chromocenters in *h2a.w-2* mutants, confirming that H2A.W regulates chromocenter formation (Supplementary Fig. 5a, b). Similarly, protein coding genes upregulated in *h2a.w-2* had diminished transcript levels in mutants complemented with a H2A.W.6 genomic construct (Supplementary Fig. 5c).

**Figure 3.**
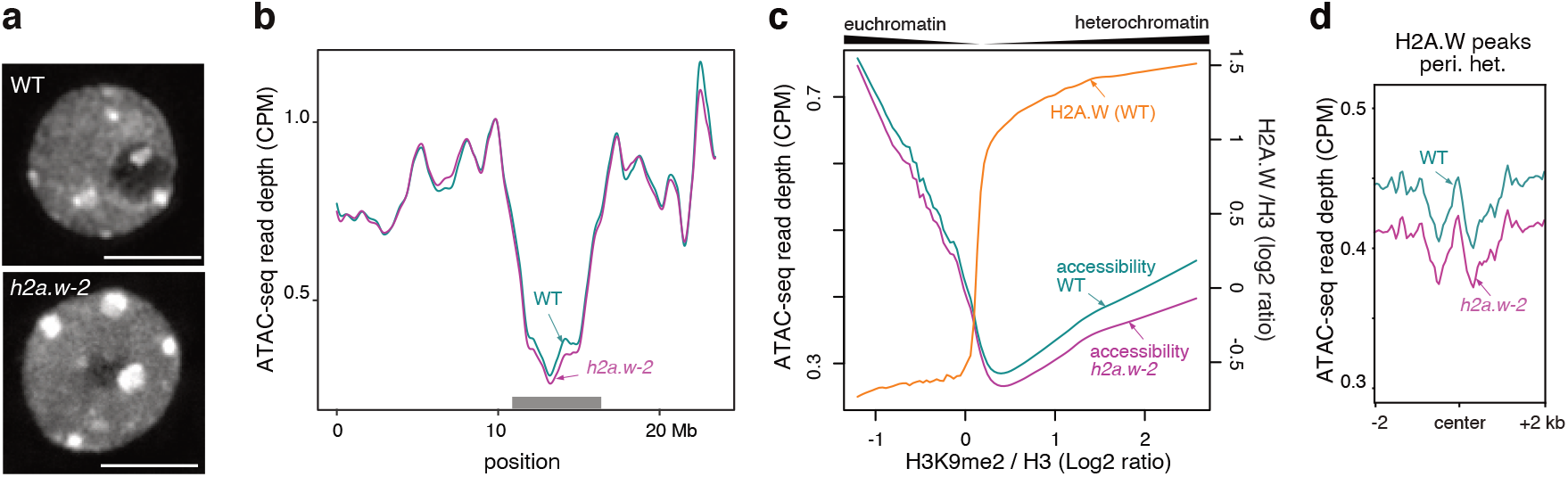
Loss of H2A.W alters heterochromatin organization and accessibility. (**a**) Representative images of DAPI-stained WT (Col-0) and *h2a.w-2* leaf interphase nuclei. Scale bar is 5 μm. (**b**) Locally weighted scatterplot smoothing (LOESS) fit of ATAC-seq read depth averaged in 1 kb bins across chromosome 3 in WT and *h2a.w-2*. Average of two replicates shown. The grey rectangle indicates the location of pericentromeric heterochromatin. (**c**) Smoothing spline fit (50 degrees of freedom) of WT H2A.W levels (log_2_ ChIP-seq H2A.W/H3) and of ATAC-seq read depth (CPM normalized) in WT and *h2a.w-2* in 1 kb windows plotted against WT H3K9me2 level. (**d**) ATAC-seq read depth over H2A.W peaks in pericentromeric heterochromatin. Average of two replicates shown.

To investigate the impact of H2A.W on chromatin accessibility, we applied the Assay for Transposase Accessible Chromatin using sequencing (ATAC-seq)^34^ to *h2a.w-2* and WT ten-day-old seedlings. In WT, the accessibility of pericentromeric chromatin and regions associated with H2A.W was low relative to other regions, as expected (Fig. 3b-d, Supplementary Fig. 6a, b). Interestingly, in heterochromatic regions (those with high levels of H2A.W and H3K9me2), chromatin accessibility was positively correlated with levels of H2A.W and H3K9me2, so that highly heterochromatic sequences were relatively more accessible than less heterochromatic sequences (Fig. 3c). Unexpectedly, we observed a modest reduction in heterochromatin accessibility in *h2a.w-2* that correlated well with WT H2A.W levels, suggesting H2A.W provides heterochromatin with a certain degree of accessibility in the WT (Fig. 3b-d and Supplementary Fig. 6a, b). This observation was at odds with previous reports showing that H2A.W promotes nucleosome thermostability^28^ and compaction of nucleosome arrays^24^, suggesting that the decreased accessibility observed in *h2a.w-2* heterochromatin may not be the direct consequence of the loss of H2A.W. We hypothesized that the reduction of chromatin accessibility in *h2a.w-2* results from the replacement of H2A.W by other types of H2A variants and/or the deposition of a chromatin component that impedes chromatin accessibility.

### H2A.X and replicative H2A replace H2A.W in *h2a.w-2* heterochromatin

To assess the composition of chromatin in *h2a.w-2,* we profiled the genome-wide distribution of H3 and other H2A variants by chromatin immunoprecipitation followed by high-throughput sequencing (ChIP-seq). Profiles of H3 enrichment determined by ChIP-seq were similar in WT and *h2a.w-2* plants, suggesting that nucleosome density was not responsible for the change in accessibility in *h2a.w-2* (Supplementary Fig. 7a). Since H2A variants confer distinct stability to nucleosomes^28^, the replacement of H2A.W by another H2A variant is expected to affect chromatin properties. Based on *in vitro* thermostability assays, replicative H2A confers higher stability than H2A.W and H2A.X, whereas H2A.Z nucleosomes are the least stable^28^. Similarly, different H2A variants are more or less favorable to the deposition of DNA methylation^30^. Therefore, we explored the distribution of the other three H2A variants in *h2a.w-2* plants by profiling H2A.X (H2A.X.3 and H2A.X.5), H2A.Z (H2A.Z.9), and replicative H2A (H2A.1 and H2A.13) using ChIP-seq. In WT, H2A.Z, H2A.X, and H2A showed relative depletion over pericentromeric heterochromatin and at H2A.W-associated regions (Fig. 4a, b), as previously reported^24^ In *h2a.w-2,* we found a striking gain of H2A.X, and to a lesser extent replicative H2A, but not H2A.Z, over regions normally marked by H2A.W in WT (Fig. 4a-c). CG DNA methylation has been shown to exclude H2A.Z from methylated DNA, and H2A.Z and CG methylation are largely anticorrelated in WT plants^30,35^. The largely unchanged H2A.Z distribution in *h2a.w-2* is consistent with our observation that CG methylation patterns are also unaltered in *h2a.w-2* (Fig. 2a, Supplementary Fig 3a, b). Western blot analyses confirmed increased levels of replicative H2A and H2A.X in *h2a.w-2* chromatin (Supplementary Fig. 7b).

**Figure 4.**
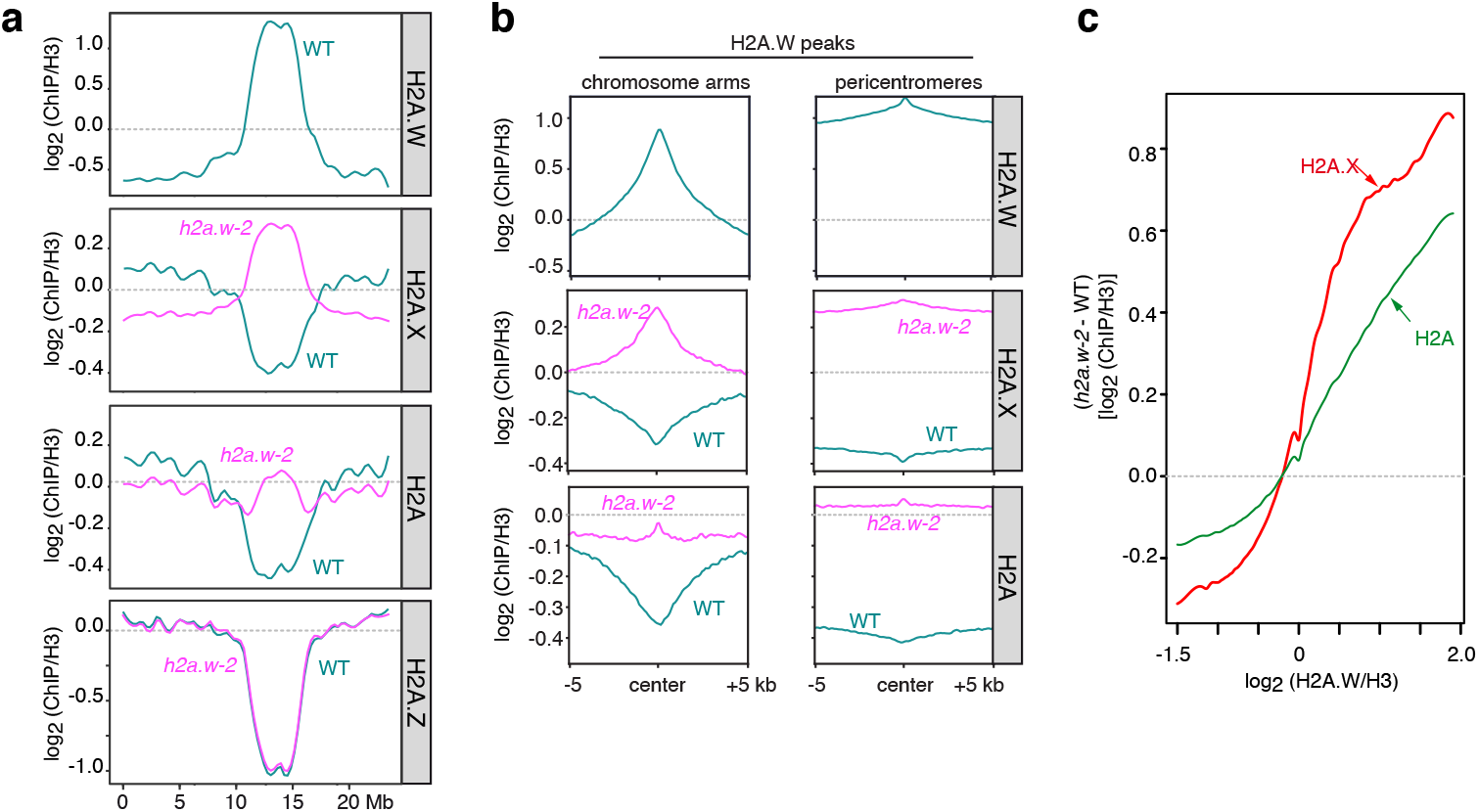
Replicative H2A and H2A.X replace H2A.W in *h2a.w-2.* (a) Locally weighted scatterplot smoothing (LOESS) fit of H2A.W, H2A.X, replicative H2A and H2A.Z levels averaged in 1 kb bins across chromosome 3 in WT and *h2a.w-2.* Average of two replicates shown. (b) Metaplots of average H2A.W, H2A.X, and replicative H2A levels from two replicates over H2A.W peaks in the chromosome arms and pericentromeres. (c) Smoothing spline fits (50 degrees of freedom) of changes in H2A.X and replicative H2A levels *(h2a.w-2* minus WT; log_2_ ChIP/H3) in 1 kb windows plotted against WT H2A.W level.

Although replicative H2A nucleosomes show higher thermal stability than H2A.W nucleosomes, *in vitro* DNA protection assays have shown that replicative H2A confers less protection than H2A.W^28^. This indicated to us that the increase in replicative H2A at pericentromeric chromatin may not be responsible for the observed decrease in accessibility. The *in vitro* thermostability of H2A.W and H2A.X nucleosomes is similar^28^ and thus also should not directly account for the changes in chromatin accessibility in *h2a.w-2.* However, H2A.X is primarily known for its role in DNA damage response, during which it becomes rapidly phosphorylated to form γH2A.X aggregates^25^, which could impact chromatin accessibility. The ratio of γH2A.X/H2A.X remained unchanged in *h2a.w-2* (Supplementary Fig. 7b), suggesting that γH2A.X was not responsible for the change in chromatin accessibility in *h2a.w-2.* In support of this conclusion, DNA damage response genes are not mis-regulated in *h2a.w-2* (Supplementary Fig. 7c).

As changes in H2A variant composition were unlikely to be directly responsible for the decreased heterochromatin accessibility in *h2a.w-2*, we next examined other epigenetic modifications. Profiles of epigenetic marks typically associated with heterochromatin, namely H3K9me1, H3K9me2, and H3K27me1, were similar in WT and *h2a.w-2* (Supplementary Fig. 7a). Hence, maintenance of these post translational modifications is independent of H2A.W and these modifications are not responsible for the change in heterochromatin accessibility in *h2a.w-2.* Intriguingly, reduced levels of non-CG methylation in *h2aw-2* were not accompanied by detectable changes in H3K9me2, suggesting that maintenance of H3K9me2 by SUVH4/5/6 may be less sensitive than CMT2 and CMT3 activities to changes in accessibility occurring in *h2a.w-2* (Supplementary Fig. 7a). Alternatively, changes in H3K9me2 could be below the detection threshold allowed by ChIP-seq.

### Heterochromatin H1 levels increase in *h2a.w-2*

Interestingly, the DNA methylation changes in *h2a.w-2* appeared to be the inverse of those previously found in linker histone H1 mutants, which display decreased DNA methylation at short euchromatic TEs and increased methylation at long heterochromatic TEs^3^. This suggested that the DNA methylation changes in *h2a.w-2* may be related to changes in H1 distribution and prompted us to explore H1 patterns in *h2a.w-2*. Consistent with earlier work^15–17^, our ChIP-seq analyses revealed that H1 is enriched in pericentromeric heterochromatin relative to euchromatin in WT (Fig. 5a). Regions associated with H2A.W in WT were also generally enriched in H1 (Fig. 5b). In *h2a.w-2,* we observed a further increase in H1 at pericentromeric heterochromatin, accompanied by a modest decrease of H1 along chromosome arms (Fig. 5a). Regions normally marked by H2A.W in both pericentromeric regions and chromosome arms also showed increased H1 enrichment in *h2a.w-2* relative to WT that correlated well with WT H2A.W levels (Fig. 5b, c), indicating that H2A.W opposes deposition of H1. Western blot analysis indicated that global nuclear H1 levels were similar in *h2a.w-2* and WT (Supplementary Fig. 7b), suggesting that the total pool of H1 available is limiting and that increased recruitment of H1 in pericentromeric heterochromatin and other H2A.W-associated regions in *h2a.w-2* is likely responsible for the relative depletion of H1 along the chromosome arms (Fig. 5a, b). Histone H1 is known to obstruct chromatin accessibility and stabilize nucleosomes by binding to the linker DNA^13^. Therefore, redistribution of H1 may account for the change in chromatin accessibility in *h2a.w-2* (Fig. 3b-d, Supplementary Fig. 6a, b).

**Figure 5.**
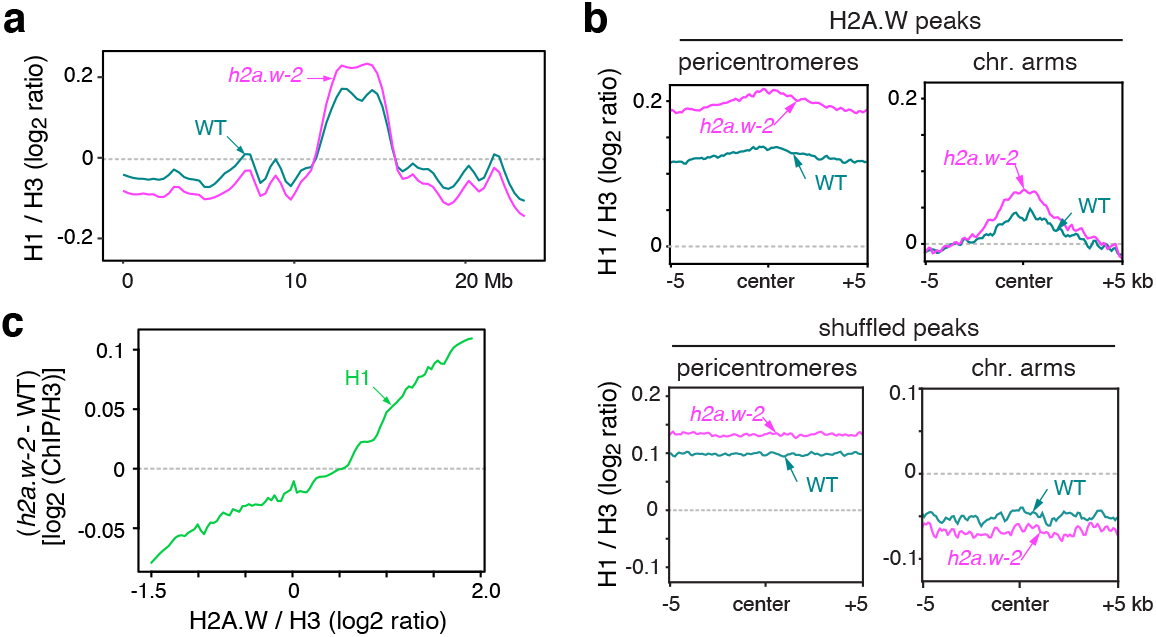
H2A.W antagonizes histone H1 deposition in heterochromatin. (a) Locally weighted scatterplot smoothing (LOESS) fit of H1 levels averaged in 1 kb bins across chromosome 3 in WT and *h2a.w-2.* Average of two replicates shown. (b) Metaplots of average H1 levels from two replicates over H2A.W peaks in pericentromeres and the chromosome arms (top panel). Plots over randomly shuffled peaks within the chromosome arms and pericentromeres are shown for comparison (bottom panel). (c) Smoothing spline fits (50 degrees of freedom) of change in H1 levels *(h2a.w-2* minus WT; log_2_ ChIP/H3) in 1 kb windows plotted against WT H2A.W level.

### H2A.W and H1 cooperatively control heterochromatin accessibility

To test whether increased H1 levels could explain the reduced accessibility and DNA methylation in *h2a.w-2* pericentromeres, we obtained *h2a.w-2 h1.1 h1.2* quintuple mutants (hereafter named *h1 h2a.w),* which lack both H2A.W and H1 (Supplementary Fig. 6c). We then compared chromatin accessibility and DNA methylation between *h2a.w-2*, *h1* and *h1 h2a.w* mutants. Chromocenters were dispersed in *h1* mutant nuclei, consistent with recent reports^15,19^, and even further dispersed in *h1 h2a.w* (Fig. 6a, b). Consistent with this, chromatin accessibility increased over pericentromeric heterochromatin regions associated with H2A.W in *h1* and further in *h1 h2a.w* (Fig. 6c, d, Supplementary Fig. 6a, b). This is in agreement with recent MNase-seq profiles showing reduced nucleosomal density in *h1* mutants^19^. Moving from chromosomal arms into pericentromeric regions, heterochromatin content and average TE length increase,^3^ and H2A.W levels are well-correlated with TE length in WT (Fig. 6e). Gain in chromatin accessibility in *h1* was also correlated with TE length and WT H2A.W levels, essentially mirroring changes in accessibility in *h2a.w-2* (Fig. 6e). Combined loss of H1 and H2A.W in *h1 h2aw* mutants led to an even stronger increase in chromatin accessibility at regions normally associated with H2A.W (Fig. 6e). This indicates that H2A.W restricts heterochromatin accessibility in the absence of H1.

**Figure 6.**
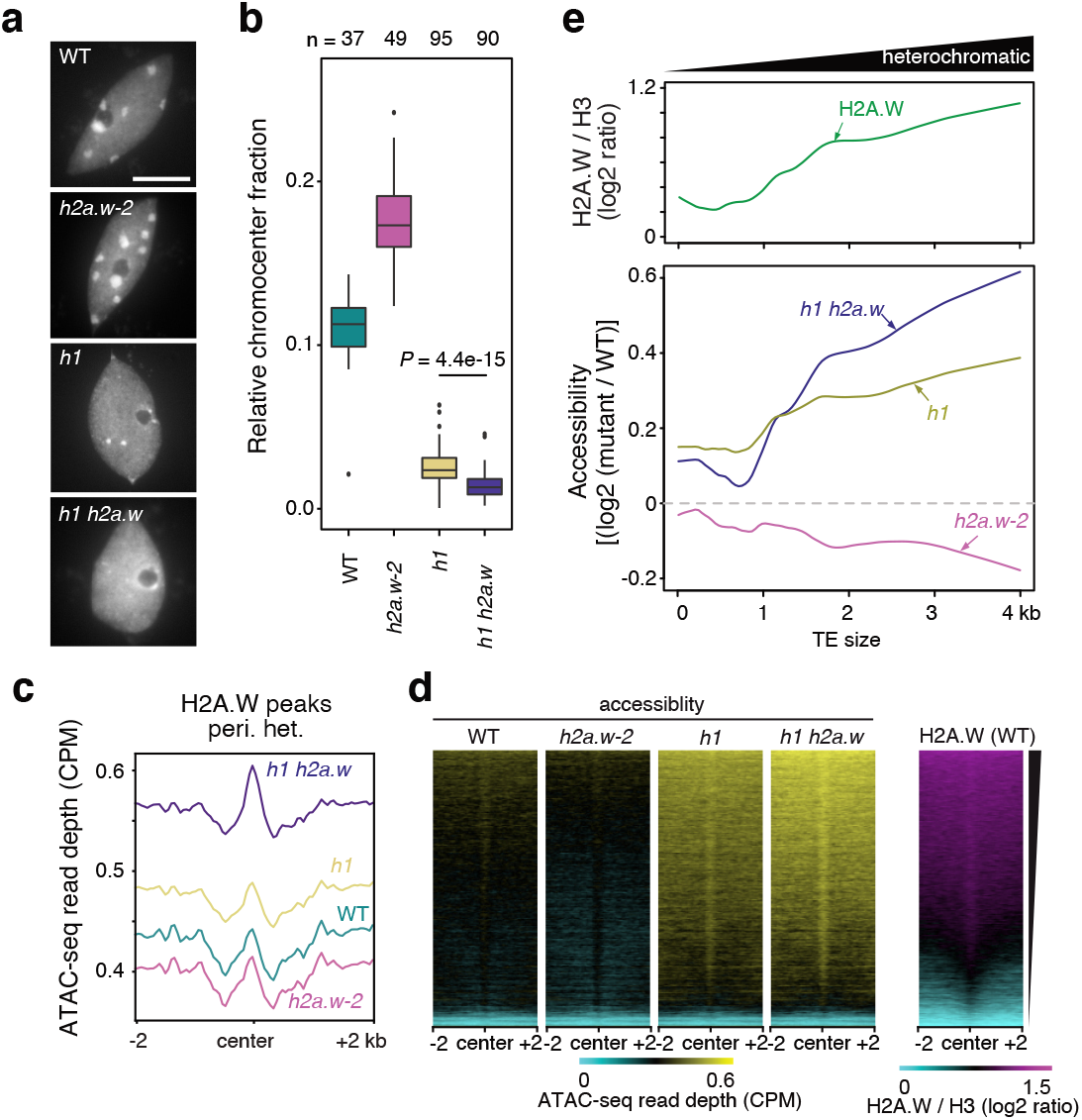
H2A.W and H1 co-regulate heterochromatin accessibility. **a** Representative images of DAPI-stained WT (Col-0), *h2a.w-2, h1* and *h1 h2a.w* leaf interphase nuclei. Scale bar is 5 μm. **b** Quantification of the relative chromocenter fraction in WT (Col-0), *h2a.w-2, h1* and *h1 h2a.w* nuclei. Number of analyzed nuclei are indicated on the top. Whiskers indicate 1.5X IQR. Outliers are represented by circles. Relative chromocenter fraction in *h1* and *h1h2a.w* show statistically significant difference (*P* = 4.4e-15, Wilcoxon rank sum test) **c** ATAC-seq read depth over H2A.W peaks in pericentromeric heterochromatin. Average of two replicates shown. **d** Heat maps of average ATAC-seq read depth over H2A.W peaks in pericentromeres ranked based on level H2A.W signal in WT **e** Locally weighted scatterplot smoothing (LOESS) fit of WT H2A.W levels (log_2_ ChIP/H3; top panel) and changes in chromatin accessibility (mutant */* WT; ATAC-seq read depth) in 50 bp windows plotted against TE size.

DNA methylation profiles of *h1* mutants were consistent with previously published data, showing increased methylation over long heterochromatic TEs and reduced CHH methylation at short TEs (Fig. 7a and Supplementary Fig. 8)^20^. Similarly, non-CG methylation levels increased at CMT2-dependent regions but were reduced at DRM2-dependent regions in *h1* (Fig. 7b, c). Again, these changes were largely opposite to those in *h2a.w-2* mutants, supporting the hypothesis that H1 is responsible for altered DNA methylation in *h2a.w-2.* H1 has been shown to control DNA methylation over heterochromatin, presumably by restricting access of DNA methyltransferases to heterochromatic DNA^3^. In concordance with increased heterochromatin accessibility in *h1 h2a.w,* we found that long heterochromatic TEs and CMT2-dependent regions further gained DNA methylation in *h1 h2a.w* compared to *h1* alone (Fig. 7b). This supports a positive correlation between heterochromatin accessibility and DNA methylation levels, and suggests that H2A.W hinders access of DNA methyltransferases to heterochromatic DNA in the absence of H1.

**Figure 7.**
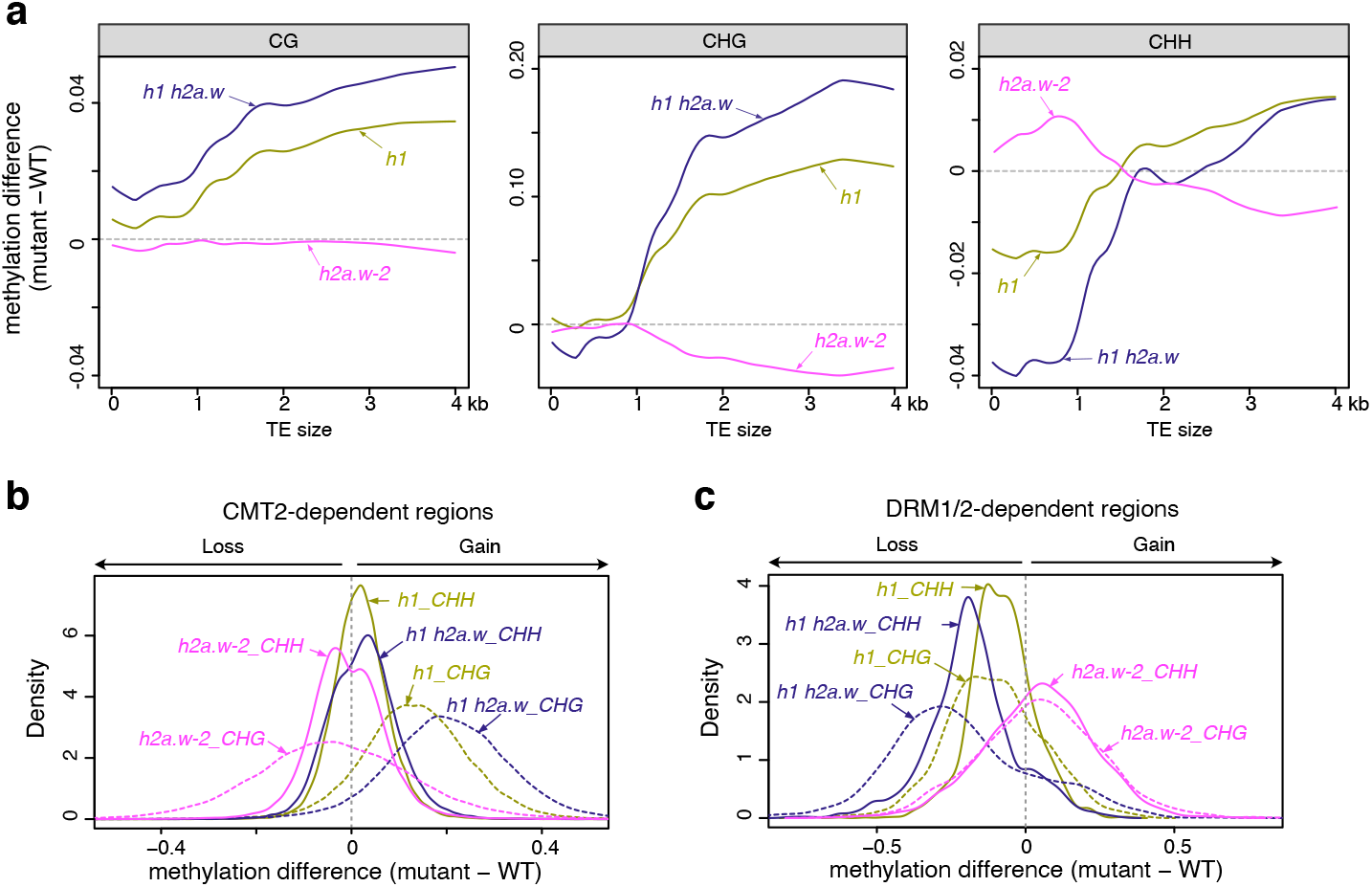
Impact of combined loss of H2A.W and H1 on DNA methylation. **a** Locally weighted scatterplot smoothing (LOESS) fit of changes in CG, CHG and CHH DNA methylation in *h2a.w-2, h1* and *h1 h2a.w* mutants (mutant minus WT) in 50 bp windows plotted against TE size. **b** Kernel density plots of CHG and CHH DNA methylation changes at CMT2-dependent regions in *h2a.w-2, h1* and *h1 h2a.w* mutants. **c** Kernel density plots of CHG and CHH DNA methylation changes at DRM1/2-dependent regions in *h2a.w-2, h1* and *h1 h2a.w* mutants.

Overall, pericentromeres gain H1 and lose both accessibility and non-CG methylation in *h2a.w-2* and these changes are strongly reversed in *h1 h2a.w* mutants (Fig. 8). This is consistent with a model wherein H2A.W blocks H1 deposition in WT. Loss of H2A.W then leads to overaccumulation of H1 at sites normally occupied by H2A.W, decreased accessibility and reduced DNA methylation.

**Figure 8.**
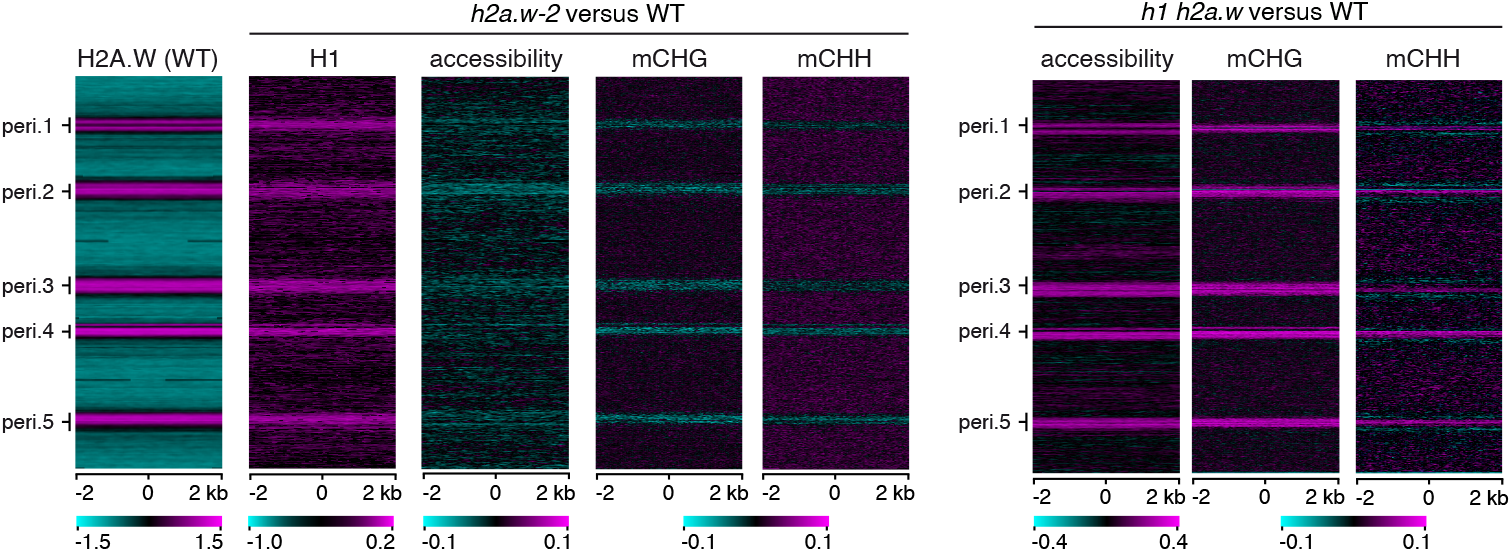
*h2a.w-2* and *h1 h2a.w* show opposite changes in accessibility and non-CG methylation. The genome was divided into consecutive, non-overlapping 4-kb bins, which are stacked according to their genomic position from the top of chromosome 1 to the bottom of chromosome 5. Pericentromeric regions are indicated (peri.1 to peri.5). H2A.W enrichment in the WT (log2 ChIP/H3) is shown as a heat map on the left. Other heat maps show changes in H1 levels (*h2a.w-2/*WT; log_2_ ChIP/H3), changes in chromatin accessibility (log2 mutant*/*WT; ATAC-seq read depth), and changes in CHG and CHH DNA methylation (mutant minus WT) in *h2a.w-2* versus WT and in *h1 h2a.w* versus WT.

## Discussion

Chromosomal rearrangements are common in T-DNA insertion lines^36,37^. Here we identified a large chromosomal rearrangement in the *hta6-1* SALK line that resulted in a duplication of the translocated region during the generation of the *hta6-1 hta7 hta12* triple mutants *(h2a.w-1).* This chromosome rearrangement, and not the loss of H2A.W, is responsible for the developmental defects and CHG hypermethylation previously reported in *h2a.w-1,* as well as the lethality of *h2a.w-1 cmt3* quadruple mutants^24^ The absence of these defects in our newly generated triple mutant, *h2a.w-2,* has now enabled us to analyze the direct impact of H2A.W on heterochromatin composition and accessibility.

H2A.W promotes higher order chromatin compaction^24^ and increases nucleosome stability^28^. Hence, we expected loss of H2A.W to increase heterochromatin accessibility. Surprisingly, we observed the opposite, with accessibility decreasing in *h2a.w-2* in association with increased recruitment of H2A, H2A.X and H1 to heterochromatin, indicating that H2A.W presence favors heterochromatin accessibility. H1 is known to stabilize the wrapping of DNA around the nucleosome, promote assembly of higher order chromatin structures^38^, and influence nucleosome spacing^39,40^. In the presence of H2A.W, loss of H1 leads to increased accessibility and increased DNA methylation in heterochromatin. These phenotypes are enhanced in the absence of both H1 and H2A.W, suggesting that H2A.W also contributes to restricting heterochromatic DNA accessibility in the absence of H1. Gain of H1 in *h2a.w-2* heterochromatin correlates well with decreased accessibility in the same regions, suggesting that H2A.W also promotes accessibility in heterochromatin by restricting H1 levels. The antagonism between H1 and H2A.W may originate from a competition for linker DNA binding. The extended C-terminal tail of H2A.W interacts with linker DNA, and this interaction prevents micrococcal nuclease accessibility^28^. The H2A.W C-terminal tail is characterized by the SPKK motif^24^, which binds A/T-rich DNA in its minor groove and causes condensation^41,42^. Among Arabidopsis H2A variants, the SPKK motif is unique to H2A.W^24^ Two SPKK-like motifs, SPAK and SP(G/A)K, are also present in the C-terminal tails of *Arabidopsis* H1.1 and H1.2 (Supplementary Fig. 9). The resulting competition between H2A.W and H1 for linker DNA binding could help prevent excessive H1 accumulation in heterochromatin. Because they do not contain SPKK motifs, H2A and H2A.X, which replace H2A.W in *h2a.w-2*, would not be able to compete with H1 for linker binding. Hence, H2A.W might allow some degree of nucleosome “breathing” in compact heterochromatin, which would facilitate access of CMT3 and CMT2 to heterochromatic DNA, enabling the maintenance of DNA methylation patterns over these regions (Figure 9). Although our data support the abovementioned scenario, we cannot exclude a potential impact of H2A and H2A.X enrichment in heterochromatin on DNA methylation and accessibility.

**Figure 9.**
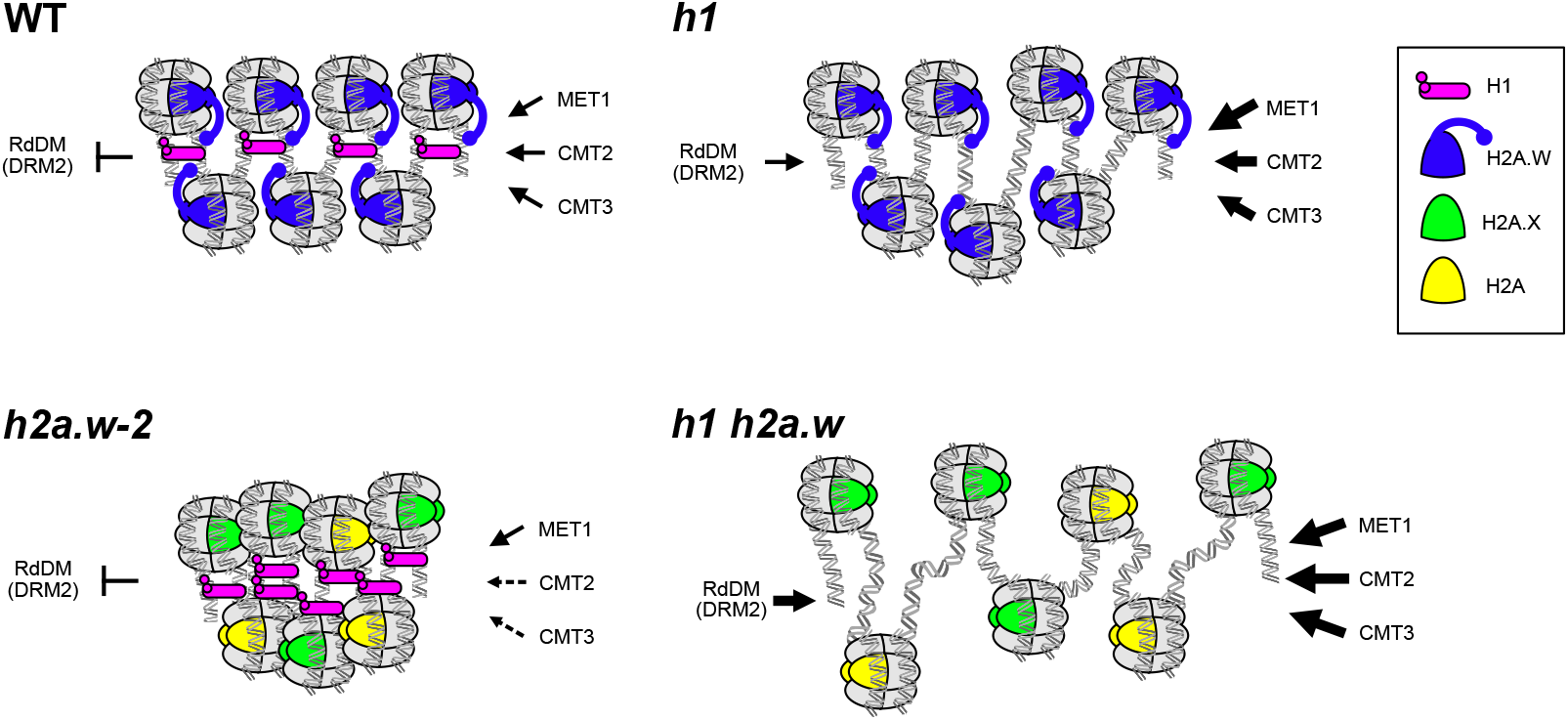
Model of co-regulation of heterochromatin accessibility by H2A.W and H1. In WT, H2A.W and H1 both interact with heterochromatin linker DNA to maintain a normal balance of accessibility to chromatin modifiers and DNA methyltransferases in heterochromatin. In the absence of H2A.W, H1 over-accumulates at linker DNA and further reduces accessibility. Loss of H1 alone causes incomplete decompaction, while loss of both H1 and H2A.W causes strong decompaction and increased accessibility. Thus, H2A.W promotes chromatin compaction in the absence of H1, but also promotes heterochromatin ‘breathing’ by opposing excessive H1 incorporation.

At genomic regions targeted by RdDM (DRM1/2-dependent regions and short TEs), which are located primarily outside of pericentromeric heterochromatin, non-CG methylation levels increase in *h2a.w-2.* These genomic regions are associated with H2A.W in WT. Because RdDM is inhibited by heterochromatin^3,32^, this suggests that H2A.W provides these regions with an heterochromatic state/identity that is lost in *h2a.w-2,* thereby facilitating RdDM-mediated DNA methylation. In agreement with a previous report, we found that DRM1/2 target regions showed reduced levels of non-CG methylation in *h1* (Fig. 7c)^3^. This loss of non-CG methylation was further aggravated in *h1 h2aw-2* (Fig. 7c). It was proposed that in the absence of H1 these RdDM-dependent genomic regions become more accessible to enzymes that catalyze euchromatic histone modifications and antagonize DNA methylation^3^. We found that chromatin accessibility at these regions was similar in *h1* and WT (Supplementary Fig. 10a). Furthermore, while DRM1/2-dependent regions are enriched in H2A.W, they are rather depleted in H1 (Supplementary Fig. 10b). Therefore, loss of H1 likely impacts DNA methylation at RdDM-dependent regions indirectly. We propose that increased heterochromatin accessibility in *h1* allows RdDM to function efficiently in heterochromatin, thus depleting the RdDM machinery from its regular targets in chromosome arms. This interpretation is consistent with the recent report that 24-nucleotide sRNAs increase in heterochromatin but decrease at euchromatic TEs in *h1* mutants^43^. As heterochromatin accessibility further increases when both H1 and H2A.W are lost, this results in a further loss of DNA methylation at usual RdDM targets in chromosome arms in *h1 h2aw-2*. Thus, maintenance of proper heterochromatin stability is also presumably important to restrain RdDM activity to specific genomic regions.

Although heterochromatin was long believed to be highly compact and inaccessible to transcriptional machinery, there is increasing evidence that low levels of accessibility within heterochromatin are required for proper heterochromatin formation by permitting access to various factors, including DNA and histone methyltransferases, that help maintain a heterochromatic state^44,45^. H2A.W is subject to specific modifications^46^, and its dynamic deposition likely participates in the regulation of chromatin accessibility through its interaction with H1 and other yet unknown factors.

## Methods

### Plant material

The *hta6* (SALK_024544C), *hta7* (GABI_149G05.01), *hta12* (SAIL_667_D09), and *cmt3-11* (SALK_148381) mutant lines used in this study were all in the Col-0 genetic background. Plants were grown in long-day conditions (16 h light, 8 h dark) at 23°C with 50% relative humidity.

### CRISPR-Cas9 targeted mutagenesis

Design of optimal guide RNA (gRNA) sequences was performed using an online bioinformatic tool (https://www.genome.arizona.edu/crispr/index.html). The spacer (GTTTCGAAATCGATGAAAGC) was ligated between the two *BbsI* sites of the pEn-Chimera vector using annealed oligonucleotides (Supplementary Table 2) and then transferred by a single site Gateway LR reaction into the pDE-CAS9 binary vector. The detailed procedure and vectors are described in^47^. Col-0 plants were transformed by floral dipping^48^ and T1 transformants were isolated following BASTA selection. Identification of heritable targeted mutagenesis events was done by PCR amplification and sequencing of the region of interest. Two independent T2 lines were then selected that had segregated away the T-DNA coding for the gRNA and Cas9 expression cassette and contained a potential insertion of a single guanine 3 bp upstream of the protospacer adjacent motif (PAM) at the gRNA-targeted HTA6 5’ coding region. The +1G insertion induces an early frame shift 89 bp downstream from the translation initiation site and a stop codon 195 bp downstream from the translation initiation site. Segregation of the mutant allele was analyzed in the T3 generation and we confirmed that both T2 lines were homozygous for the mutation. The knockout nature of this *hta6-2* allele was confirmed by immunoblot analysis using a specific antibody (see Fig. 1d). In subsequent crosses, a dCAPS assay was used to identify the *hta6-2* allele through a single *Bsa*BI digestion (cuts the mutant allele) of the PCR product (Supplementary Table 2).

### Southern blot

Genomic DNA was extracted from rosette leaves using the Wizard^®^ Genomic DNA Purification Kit (Promega) following manufacturer’s instructions. 750 ng of DNA was digested overnight with 20 units of high fidelity *SspI* restriction enzyme (New England Biolabs) in Cutsmart^®^ buffer and electrophoresed through a 1% agarose (w/v) gel for 8 h. The gel was depurinated (10 min in 0.25 N HCl), rinsed, denatured (30 min in 0.4 N NaOH, 1.5 M NaCl), neutralized (30 min in 0.5 M Tris-HCl, 1.5 M NaCl) and capillary blotted onto a Hybond-N+ membrane (Amersham) overnight. The membrane was UV-crosslinked at 150 mJ. The DNA probe was amplified from Col-0 DNA with primers indicated in Supplementary Table 2, gel-purified, and labeled with a-^32^P-dCTP using the random hexamer priming method (Megaprime DNA labeling system; Amersham) following manufacturer’s instructions and subsequently purified on illustra MicroSpin S-200 HR columns (GE Healthcare Life Sciences). Hybridization was performed using the PerfectHyb™ Plus hybridization buffer (Sigma) following manufacturer’s instructions, with overnight hybridization at 65°C followed by one washing step (10 min) in 2X SSC 0.1% SDS and two washing steps (15 min each) in 0.5X SSC 0.1% SDS, all at 65°C. The membrane was imaged on a Typhoon FLA 7000 (GE Healthcare Life Sciences).

### Inverse PCR (iPCR)

250 ng of Col-0 and *hta6-1* genomic DNA were digested by *Ssp*I and then column-purified using the Neo Biotech gel extraction kit. To favor self-recircularization of the *Ssp*I-digested fragments, ligation was performed at 15°C for 16 hours using 100 ng of the digested DNAs and 4.5 U of T4 DNA ligase (Promega) in a final volume of 100 μl. Following column purification (Neo Biotech gel extraction kit), 1/50 of the eluted DNA was used as a template for a first round of iPCR with primers that closely match the expected extremity of the translocation. A second round of PCR amplification was done using nested primers and a 1/100th dilution of the first amplification as template. A specific product of around 2.1 kb was obtained for the *hta6-1* genomic DNA and sequenced (Eurofins) to identify the left border of the translocation. The primers used for iPCR are reported in Supplementary Table 2.

### Transcript analysis

Total RNA was extracted with TRI Reagent (Sigma^®^) from 30 to 40 mg of fresh material following the manufacturer’s instructions. 8 μg of RNA were treated for 1 h at 37°C with 12 units of RQ1 DNase (Promega^®^) followed by phenol-chloroform extraction and ethanol precipitation of RNA which was subsequently dissolved in water. One-step reverse-transcription quantitative PCR (RT-qPCR) was performed with the SensiFAST™ SYBR^®^ No-ROX One-Step kit (Bioline^®^) on an Eco™ Real-Time PCR System (Ilumina^®^) with the following program: 10 min at 45°C, 5 min at 95°C, and 40 cycles of 20 s at 95°C and 30 s at 60°C. A melting curve was generated at the end of the program to control for amplification specificity. Data was normalized to a reference gene and analyzed according to the 2^_AACt^ method. Means and standard errors of the mean were calculated from independent biological samples. Differences in the means for RT-qPCR data were tested using an unpaired Student’s t-test with Welch’s correction with the *t.test* function of R version 3.4.0^49^.

### Nuclear protein extraction and immunoblot

Nuclear protein extracts for Western blot analyses were prepared as described in^25^ with few modifications. For each sample 300mg of 10-day old seedlings or 200 mg of floral buds (for H2A.W antibody characterization) are frozen in liquid nitrogen and disrupted in 2ml Eppendorf tubes using Qiagen TissueLyser II and metal beads to fine powder. Total ground powder is transferred into 15ml falcon tube containing 5ml of nuclei isolation buffer (NIB; 10 mM MES-KOH pH 5.3, 10 mM NaCl, 10 mM KCl, 250 mM sucrose, 2.5 mM EDTA, 2.5 mM β-mercaptoethanol, 0.1 mM spermine, 0.1 mM spermidine, 0.3% Triton X-100) and supplemented with protease and phosphatase inhibitors (Roche), followed by vortexing until a fine suspension was obtained. The suspension was filtered through two layers of Miracloth into 50 ml Falcon tubes, followed by washing the Miracloth with 10 ml of NIB. Remaining buffer was carefully squeezed out of the Miracloth into the tube. Nuclei were pelleted by centrifugation at 3,000 rpm at 4°C for 5 min. The pellet was washed once with 5 ml of NIB and centrifuged again. Nuclei were re-suspended in 1 ml of NIB and transferred to Eppendorf tubes followed by centrifugation for 5 min at 4°C at maximum speed. Finally, nuclei were re-suspended in 150 μl of 1x PBS supplemented with protease and phosphatase inhibitors (Roche), mixed with 50 μl of 4x Laemmli loading buffer and boiled for 5 minutes. Once the samples reached room temperature, 2 μl of Benzonase (Millipore) was added and incubated for 10min on bench. Samples are again boiled for 3min to inactivate Benzonase. Samples were spun at maximum speed for 5 min to pellet down insoluble fraction and supernatant is transferred to fresh Eppendorf tubes. For Western blot analyses, 10 μl for histone variants and 5 μl for H3 (used as a loading control) were loaded per lane. Nuclear proteins were resolved using NuPAGE 4-12% Bis-Tris protein gels. Resolved proteins were transferred onto PVDF membrane using Bio-Rad wet transfer unit. Western blot analysis was performed using 1:1000 diluted antibodies in 5% milk in TBST. H2A.W.6, H2A, H2A.X and H2A.Z antibodies are affinity-purified rabbit polyclonal antibodies made by GenScript USA Inc (Piscataway, NJ) against peptides GGRKPPGAPKTKSVC, CPKKAGASKPSADE, CKVGKNKGDIGSASQ, and KPSGSDKDKDKKKPC, respectively, as described previously^24^. H2A.W.7, and γH2A.X antibodies^25^ were described previously. H2A.W and H1 antibodies were generated using peptides CTTKTPKSPSKATKSP and CRTGSSQYAIQKFIEEK respectively at Eurogentec.

### Whole Genome Bisulfite sequencing (BS-seq)

Genomic DNA was extracted from the aerial portions of 10-day old seedlings using the Wizard^®^ Genomic DNA Purification Kit (Promega) following manufacturer’s instructions. For WT and *h2a.w-2* replicates 1 and 2, sodium bisulfite conversion, library preparation, and sequencing on a Hiseq 4000 were performed at the Beijing Genomics Institute (Hong Kong) from 1 μg DNA, producing paired 100-bp (replicate 1) or 150-bp (replicate 2) paired-end reads. For remaining samples, methylated adapters were ligated prior to bisulfite treatment using the ZYMO EZ DNA methylation Gold kit, libraries were prepared using the Nugen ultralow methyl-seq kit, and sequenced on NovaSeq 6000. Reads were trimmed, mapped to the TAIR10 genome, and methylation called using methylpy (version 1.4.3). Only uniquely mapping reads were retained. Pericentromeres and chromosome arm regions were defined based on H3K9me2 distribution^50^.

### RNA-seq

Total RNA was isolated with RNeasy Mini kit (Qiagen) from 10-day old seedlings in three replicates. DNase treatment was done on 2 μg of total RNA with DNA free DNase Kit (Invitrogen). From 1 μg of total RNA, rRNA was depleted using RiboZero kit (Illumina). NGS-libraries were generated using NEBnext Ultra II directional RNA library prep kit for Illumina and sequenced as PE75 reads on an Illumina NextSeq550.

Reads were trimmed and filtered for quality and adapter contamination using Trim Galore^51^ and aligned to the TAIR10 genome using STAR^52^. Reads aligning equally well to more than one position in the reference genome were discarded, and probable PCR duplicates were removed using MarkDuplicates from the Picard Tools suite^53^. Alignment statistics for each library are available in Supplementary Table 3. Read counts for each gene and TE were obtained using htseq-count^54^, with annotations from araport11^55^. Annotated TEs overlapping strongly (> 80%) with an annotated TE gene were considered TE genes, and the TE annotation was discarded. Differential expression analysis was performed using DESeq2^56^, and genes were considered differentially expressed with an adjusted *p*-value < 0.05 and abs[log_2_(fold change)] > 1.

### ATAC-seq

ATAC-seq was performed as described in^57^. Briefly, 0.5 g of freshly collected 10-day old seedlings were chopped in 4 ml of pre-chilled lysis buffer (15 mM Tris-HCl pH 7.5, 20 mM NaCl, 80 mM KCl, 0.5 mM spermine, 5mM B-mercaptoethanol, 0.2% Triton X-100). After chopping, the suspension was filtered through a 40 μM filter. Nuclei were further enriched using a sucrose gradient. Enriched nuclei were resuspended in 0.5 ml of pre-cooled lysis buffer with 4,6-Diamidino-2-Phenylindole (DAPI) and incubated for 15 min. DAPI stained nuclei were sorted on FACS Aria III (BD Biosciences). Sorted nuclei (50,000) were pelleted and washed once (10 mM Tris-HCl pH 8.0, 5 mM MgCl2). Tagmentation reaction was carried out using Nextera reagents (TDE1 Tagment DNA Enzyme (Catalog No. 15027865), TD Tagment DNA Buffer (Catalog No. 15027866)). Tagmented DNA was isolated using Qiagen MinElute PCR purification kit. NGS libraries were amplified using NEBNext high fidelity 2X master mix and Nextera primers. The number of PCR cycles was determined using a method described in^58^. NGS-libraries were PE75 sequenced on an Illumina NextSeq550.

Reads were trimmed and filtered as indicated above (RNA sequencing), and aligned to TAIR10 using bowtie2^59^. Reads aligning to multiple positions and PCR duplicates were removed (see RNA sequencing). Only properly paired reads were retained for the analysis. Alignment statistics for each library are available in Supplementary Table 3. Sample tracks and peaks in WT and *h2a.w-2* were obtained using Genrich^60^ with parameters -p 0.01, -a 200, -l 100 and -g 100. ChrM, ChrC, and several rRNA regions with very high coverage were omitted from the analysis. Metaplots of ATAC-seq signal over various genomic regions were created using deeptools^61^. Plots of ATAC-seq signal over entire chromosomes are based on average signal over 1 kb non-overlapping bins tiled genome-wide, calculated using deeptools. Smoothed conditional mean of the signal was computed using the LOESS smoothing method with bin width span 0.1 and plotted using R^49^. A comparison of the ATAC-seq data generated in this study with selected published ATAC-seq datasets is provided in Supplementary Fig. 11.

### ChIP-seq

ChIP was performed as described in^62^. Briefly, 3 g (approx. 0.3 mg for each immunoprecipitation (IP)) of 10-day old seedlings were fixed in 1% PFA. Fixed seedlings were ground to fine powder in liquid nitrogen using a mortar and pestle. Nuclei were isolated using M2 buffer (10 mM phosphate buffer pH 7.0, 100 mM NaCl, 10 mM β-mercaptoethanol, 10 mM MgCl_2_, 0.5% Triton X-100, 1 M hexylene glycol, 1* cOmplete protease inhibitor cocktail) and M3 buffer (10 mM phosphate buffer pH 7.0, 100 mM NaCl, 10 mM B-mercaptoethanol, 1* cOmplete protease inhibitor cocktail). Chromatin shearing was done using a Covaris E220 with the following settings: treatment time 15 minutes, acoustic duty factor % 5.0, PIP 140 W, Cycles per burst 200 and max temperature 8°C. IP, washes, and DNA isolation were carried out as described in^62^. 5 μg of the following antibodies were used for IP: H2A, H2A.X, H2A.Z (described in^24^). H3 (ab1791 Abcam), H3K9me1 (ab8896 Abcam), H3K9me2 (ab1220/Abcam), H3K27me1 (17-643/Millipore) and H1 (AS111801/Agrisera) antibodies were obtained from commercial sources. NGS libraries were generated using Ovation Ultralow Library System V2 (NuGEN) for replicate 1 and NEBNext Ultra II DNA preparation kit for replicate 2. NGS-libraries were SR75 sequenced on an Illumina NextSeq550.

Reads were trimmed, filtered, and aligned using bowtie2, and multi-mapping reads and PCR duplicates were removed, all as indicated above (see **ATAC-seq**). Alignment statistics for each library are available in Supplementary Table 3. Sample tracks and metaplots over genomic regions were obtained using deeptools^61^ bamCoverage (--normalizeUsing CPM). All samples except for H3 were normalized to their matched H3 sample using deeptools bamCompare. Plots over entire chromosomes were obtained from average ChIP-seq signal over 1 kb non-overlapping bins tiled genome-wide and smoothed using the same approach as the ATAC-seq data. H2A.W ChIP-seq data were re-analyzed from^24^.

## Data availability

The data supporting the findings of this study are available within the article and its Supplementary Information. High throughput sequencing data has been deposited in the Gene Expression Omnibus (GEO) database and can be accessed with the accession number GSE146948. All data are available from the corresponding author upon reasonable request.

## Acknowledgments

We thank James Watson from GMI for editing and comments on the manuscript, and we thank the Vienna Biocenter Core Facility Next Generation Sequencing. Work in the Mathieu laboratory was supported by CNRS, Inserm, Université Clermont Auvergne, Young Researcher grants from the Auvergne Regional Council (to O.M.), an EMBO Young Investigator award (to O.M.), and a grant from the European Research Council (ERC, I2ST 260742 to O.M.). P.B. was supported by a PhD studentship from the Ministère de l’éducation nationale, de l’enseignement supérieur et de la recherche. Work in the Berger laboratory was supported by the Gregor Mendel Institute core funding from the Austrian Academy of Sciences and the Austrian Science Fund (FWF): I2303, P32054, P28320, and P26887. Work in the Jacobsen laboratory was supported by the National Institutes of Health (R35GM130272 to S.E.J.). C.L.P was supported by the National Institutes of Health under a Ruth L. Kirschstein National Research Service Award (F32GM136115). S.E.J. is an Investigator of the Howard Hughes Medical Institute. The funders had no role in study design, data collection and analysis, decision to publish, or preparation of the manuscript.

## Author contributions

P.B., C.L.P., R.Y., F.B, S.E.J. and O.M. designed the study. P.B., R.Y., T.P., Z.J.L. and M.N.P. performed experiments. C.L.P. and O.M performed bioinformatic analyses. P.B., C.L.P., R.Y., F.B. and O.M. wrote the manuscript. F.B., S.E.J. and O.M. coordinated the research.

---

**Supplementary Fig. 1.** (a) Sequencing coverage of published *h2a.w-1* BS-seq and RNA-seq data^24^. Locally weighted scatterplot smoothing (LOESS) fit of the log2 (*h2a.w-1* / WT) coverage ratio over non-overlapping 1 kb windows along the five *Arabidopsis* chromosomes. The black arrow indicates the genomic location of *CMT3.* (b) Scheme of a backcross between WT (Col-0) and *hta6-1.* Schematic representation of chromosomes 1 and 5 in the parental plants is shown. The *hta6-1* line harbors a large translocation of chromosome 1 to chromosome 5 linked to the *HTA6* gene (in red). A schematic representation of the T-DNA-associated rearrangements is shown. Partial characterization of these rearrangements was done by inverse PCR on *hta6-1*-circularized DNA after *Ssp*I digestion and by direct PCR amplification using primers specific to chromosome 1 and 5 and a T-DNA-specific primer (black arrows). On chromosome 5, the orientation of the chromosome 1 translation is unknown and is shown arbitrarily. The translocated part is flanked by T-DNA sequences at both sides in a likely complex genomic structure as we were unable to PCR amplify chromosome 1 / chromosome 5 junctions. At chromosome 1, the ~5 Mb missing region is replaced by T-DNA/vector sequences. Plant pictures show representative images of the three plant phenotypes segregating in the F2 progeny of the Col-0 x *hta6-1* cross (scale bar = 1 cm). PCR analysis of the segregation of the *hta6-1* T-DNA and the chromosome 1 and 5 rearrangements in F2 plants of each phenotype is shown. Genomic dose of the chromosome 1 translocated region is indicated below the gels, together with a schematic representation of the inferred corresponding chromosome 1 and 5 structures. (c) Western blot confirming the specificity of the H2A.W antibody. The antibody was tested on total nuclear extracts. No band is detected in *h2a.w-2* suggesting its specificity to H2A.W. Furthermore, the H2A.W antibody reacts to all H2A.W-RFP fusion proteins (bands indicated by the arrow) with different affinities, indicating that the antibody recognizes all three H2A.W variants. H3 antibody staining is carried out as loading control. Bands marked by an asterisk indicate a degradation product of H2A.W-RFP.

**Supplementary Fig. 2.** (a) MA-plots of genes and TEs obtained using DEseq2^56^. Genes were separated into protein coding, transposable element genes, and all other genes (ncRNAs, etc.). Significantly upregulated genes in *h2a.w-2* are highlighted in red, downregulated genes are in blue. (b) Summary of differentially expressed genes and TEs in *h2a.w-2.* (c) Quantification of *CMT3* transcripts by RT-qPCR in the indicated genotypes. L/M and S refer to Large/Medium and Small plant phenotypes, respectively (see Supplementary Fig. 1b). Plants with an S phenotype contain four doses of the chromosome 1 rearranged region. Statistically significant differences between the means from mutant and WT were tested with an unpaired two-sided Student’s t-test (*: *P* < 0.05). CMT3 transcript levels are normalized to *ACT2* and further normalized to Col-0. Sample means are shown with error bars representing standard error of the mean (n=3 for *hta6-1* F3 BC samples, n=4 for others). (d) The *h2a.w-2 cmt3* quadruple mutants develop normally. Representative pictures of WT and *h2a.w-2 cmt3* plants.

**Supplementary Fig. 3. (a)** Average DNA methylation levels in the CG, CHG, and CHH sequence contexts, in 100 kb windows across chromosome 4 in individual WT and *h2a.w-2* replicates. (b) CG, CHG, and CHH methylation levels over all Arabidopsis protein coding genes (PCGs) and all Arabidopsis TEs in WT and *h2a.w-2*. PCGs and TEs were aligned at the 5’ (left dashed line) and 3’ end (right dashed line), and sequences 3 kb upstream or downstream were included, respectively. Average methylation over 100 bp bins is plotted. (c) Metaplots of CHG and CHH methylation levels over TEs located in pericentromeric heterochromatin and TEs located in chromosome arms in WT and *h2a.w-2*. (d) CHG and CHH methylation levels over H2A.W peaks in the chromosome arms and pericentromeres.

**Supplementary Fig. 4.** Flow cytometry profiles of nuclei isolated from 10-day old seedlings of Col-0 and *h2a.w-2.* The X-axis represents ploidy level or DNA content and the Y-axis represents the number of nuclei.

**Supplementary Fig. 5.** (a) Western blot confirming expression of H2A.W.6 in three *h2a.w-2* complementation lines and WT and *h2a.w-2* controls. (b) Partial rescue of relative chromocenter fraction in H2A.W.6-complemented *h2a.w-2* lines. Number of analyzed nuclei are indicated on the top. Whiskers indicate 1.5X IQR. Outliers are represented by circles. Relative chromocenter fraction in rescued lines significantly lower than in *h2a.w-2 (P* = 6.1e-4, Wilcoxon rank sum test). (c) Relative expression of three genes upregulated in *h2a.w-2* in WT, *h2a.w-2,* and *h2a.w-2*::H2A.W.6 by qPCR.

**Supplementary Fig. 6.** (a) Locally weighted scatterplot smoothing (LOESS) fit of ATAC-seq read depth averaged in 50 kb bins across all *Arabidopsis* chromosomes in indicated genotypes. Average of two replicates is shown. (b) Genome browser views of genomic regions on the left arm (top) and on the pericentromere (bottom) of chromosome 3. The two ATAC-seq replicates for each genotype are shown. (c) Western blot confirming *h1 h2a.w* quintuple loss of function mutant

**Supplementary Fig. 7.** (a) Locally weighted scatterplot smoothing (LOESS) fit of H3, H3K9me1, H3K9me2, and H3K27me1 levels averaged in 1 kb bins across chromosome 3 in WT and *h2a.w-2*. Average of two replicates shown. (b) Western blot comparing replicative H2A, H2A.X, γH2A.X, H2A.Z, and H1 protein levels between Col-0 and *h2a.w-2.* Western was performed on nuclear extracts. H3 is used as a loading control. Bar plot represents the quantification of protein levels. The plot was generated using data from two independent experiments. (c) Comparison of the expression levels of 13 genes upregulated in response to DNA damage^63^ in *h2a.w-2* and WT.

**Supplementary Fig. 8.** CG, CHG, and CHH methylation levels over TEs in pericentromeres and chromosome arms in WT, *h1* and *h1 h2a.w.* TEs were aligned at the 5’ (left dashed line) and 3’ end (right dashed line), and sequences 3 kb upstream or downstream were included, respectively. Average methylation over 100 bp bins is plotted.

**Supplementary Fig. 9.** Alignment of *Arabidopsis* Histone H1.1 and H1.2 generated using Clustal Omega (https://www.ebi.ac.uk/Tools/msa/clustalo/). SPKK-like motifs in C-terminal tails of H1 are highlighted in blue.

**Supplementary Fig. 10. (a)** ATAC-seq read depth over DRM1/2- and CMT2-dependent regions in the indicated genotypes. Average of two replicates shown. (**b**) Wild-type levels of H2A.W and H1 over DRM1/2- and CMT2-dependent regions.

**Supplementary Fig. 11. (a)** Hierarchical heatmap of ATAC-seq replicates described in this study (WT_1 and WT_2) and published ATAC-seq datasets based on Spearman correlation coefficients. Clusters were constructed using complete linkage. Colors represent the correlation coefficients that are also indicated in each box. **(b)** Genome browser views of H2A.Z ChIP-seq replicates described in this study and H2A.Z ChIP-seq replicates published in Potok et al 2019^64^

**Supplementary Table 1.** List of TEs with significantly increased copy number in *h2a.w-1* vs. WT BS-seq data^24^

**Supplementary Table 2.** List of primers used in this study.

**Supplementary Table 3.** Next-generation sequencing reads – total read counts and mapping statistics.

## References

1. Du, J., Johnson, L. M., Jacobsen, S. E. & Patel, D. J. DNA methylation pathways and their crosstalk with histone methylation. Nat. Rev. Mol. Cell Biol. 16, 519–532 (2015).

2. Stroud, H. et al. Non-CG methylation patterns shape the epigenetic landscape in Arabidopsis. Nat. Struct. Mol. Biol. 21, 64–72 (2014).

3. Zemach, A. et al. The Arabidopsis nucleosome remodeler DDM1 allows DNA methyltransferases to access H1-containing heterochromatin. Cell 153, 193–205 (2013).

4. Du, J. et al. Dual Binding of Chromomethylase Domains to H3K9me2-Containing Nucleosomes Directs DNA Methylation in Plants. Cell 151, 167–180 (2012).

5. Stroud, H. et al. Non-CG methylation patterns shape the epigenetic landscape in Arabidopsis. Nat. Struct. Mol. Biol. 21, 64–72 (2014).

6. Du, J. et al. Mechanism of DNA methylation-directed histone methylation by KRYPTONITE. Mol. Cell 55, 495–504 (2014).

7. Johnson, L. M. et al. The SRA Methyl-Cytosine-Binding Domain Links DNA and Histone Methylation. Curr. Biol. 17, 379–384 (2007).

8. Li, X. et al. Mechanistic insights into plant SUVH family H3K9 methyltransferases and their binding to context-biased non-CG DNA methylation. Proc. Natl. Acad. Sci. 115, E8793–E8802 (2018).

9. Matzke, M. A. & Mosher, R. A. RNA-directed DNA methylation: An epigenetic pathway of increasing complexity. Nat. Rev. Genet. 15, 394–408 (2014).

10. Law, J. A. et al. Polymerase IV occupancy at RNA-directed DNA methylation sites requires SHH1. Nature 498, 385–389 (2013).

11. Zhang, H. et al. DTF1 is a core component of RNA-directed DNA methylation and may assist in the recruitment of Pol IV. Proc. Natl. Acad. Sci. 110, 8290–8295 (2013).

12. Jacob, Y. et al. ATXR5 and ATXR6 are H3K27 monomethyltransferases required for chromatin structure and gene silencing. Nat. Struct. Mol. Biol. 16, 763–768 (2009).

13. Cutter, A. R. & Hayes, J. J. A brief review of nucleosome structure. FEBS Lett. 589, 2914–2922 (2015).

14. Fyodorov, D. V, Zhou, B.-R., Skoultchi, A. I. & Bai, Y. Emerging roles of linker histones in regulating chromatin structure and function. Nat. Rev. Mol. Cell Biol. 19, 192–206 (2018).

15. Choi, J., Lyons, D. B., Kim, M. Y., Moore, J. D. & Zilberman, D. DNA Methylation and Histone H1 Jointly Repress Transposable Elements and Aberrant Intragenic Transcripts. Mol. Cell 77, 310–323.e7 (2020).

16. Rutowicz, K. et al. A specialized histone H1 variant is required for adaptive responses to complex abiotic stress and related DNA methylation in Arabidopsis. Plant Physiol. pp.00493.2015 (2015). doi:10.1104/pp.15.00493

17. Wollmann, H. et al. The histone H3 variant H3.3 regulates gene body DNA methylation in Arabidopsis thaliana. Genome Biol. 18, 94 (2017).

18. He, S., Vickers, M., Zhang, J. & Feng, X. Natural depletion of histone H1 in sex cells causes DNA demethylation, heterochromatin decondensation and transposon activation. Elife 8, (2019).

19. Rutowicz, K. et al. Linker histones are fine-scale chromatin architects modulating developmental decisions in Arabidopsis. Genome Biol. 20, 157 (2019).

20. Zemach, A. et al. The arabidopsis nucleosome remodeler DDM1 allows DNA methyltransferases to access H1-containing heterochromatin. Cell 153, 193–205 (2013).

21. Lyons, D. B. & Zilberman, D. DDM1 and Lsh remodelers allow methylation of DNA wrapped in nucleosomes. Elife 6, (2017).

22. Braunschweig, U., Hogan, G. J., Pagie, L. & van Steensel, B. Histone H1 binding is inhibited by histone variant H3.3. EMBO J. 28, 3635–45 (2009).

23. Lin, C.-J., Conti, M. & Ramalho-Santos, M. Histone variant H3.3 maintains a decondensed chromatin state essential for mouse preimplantation development. Development 140, 3624–34 (2013).

24. Yelagandula, R. et al. The histone variant H2A.W defines heterochromatin and promotes chromatin condensation in arabidopsis. Cell 158, 98–109 (2014).

25. Lorkovic, Z. J. et al. Compartmentalization of DNA Damage Response between Heterochromatin and Euchromatin Is Mediated by Distinct H2A Histone Variants. Curr. Biol. 27, 1192–1199 (2017).

26. Buschbeck, M. & Hake, S. B. Variants of core histones and their roles in cell fate decisions, development and cancer. Nat. Rev. Mol. Cell Biol. 18, 299–314 (2017).

27. Lei, B. & Berger, F. H2A Variants in Arabidopsis: Versatile Regulators of Genome Activity. Plant Commun. 1, 100015 (2020).

28. Osakabe, A. et al. Histone H2A variants confer specific properties to nucleosomes and impact on chromatin accessibility. Nucleic Acids Res. 46, 7675–7685 (2018).

29. Carter, B. et al. The Chromatin Remodelers PKL and PIE1 Act in an Epigenetic Pathway That Determines H3K27me3 Homeostasis in Arabidopsis. Plant Cell 30, 1337–1352 (2018).

30. Zilberman, D., Coleman-Derr, D., Ballinger, T. & Henikoff, S. Histone H2A.Z and DNA methylation are mutually antagonistic chromatin marks. Nature 456, 125–129 (2008).

31. Zhou, W., Zhu, Y., Dong, A. & Shen, W.-H. Histone H2A/H2B chaperones: from molecules to chromatin-based functions in plant growth and development. Plant J. 83, 78–95 (2015).

32. Schoft, V. K. et al. Induction of RNA-directed DNA methylation upon decondensation of constitutive heterochromatin. EMBO Rep. 10, 1015–1021 (2009).

33. Jacob, Y. et al. Regulation of heterochromatic DNA replication by histone H3 lysine 27 methyltransferases. Nature 466, 987–991 (2010).

34. Bajic, M., Maher, K. A. & Deal, R. B. Identification of Open Chromatin Regions in Plant Genomes Using ATAC-Seq. in Methods in molecular biology (Clifton, N.J.) 1675, 183–201 (2018).

35. Coleman-Derr, D. & Zilberman, D. Deposition of Histone Variant H2A.Z within Gene Bodies Regulates Responsive Genes. PLoS Genet. 8, e1002988 (2012).

36. Clark, K. A. & Krysan, P. J. Chromosomal translocations are a common phenomenon in Arabidopsis thaliana T-DNA insertion lines. Plant J. 64, 990–1001 (2010).

37. Jupe, F. et al. The complex architecture and epigenomic impact of plant T-DNA insertions. PLoS Genet. 15, e1007819 (2019).

38. Allan, J., Mitchell, T., Harborne, N., Bohm, L. & Crane-Robinson, C. Roles of H1 domains in determining higher order chromatin structure and H1 location. J. Mol. Biol. 187, 591–601 (1986).

39. Blank, T. A. & Becker, P. B. Electrostatic Mechanism of Nucleosome Spacing. J. Mol. Biol. 252, 305–313 (1995).

40. Fan, Y. et al. H1 Linker Histones Are Essential for Mouse Development and Affect Nucleosome Spacing In Vivo. Mol. Cell. Biol. 23, 4559–4572 (2003).

41. Churchill, M. E. & Suzuki, M. ‘SPKK’ motifs prefer to bind to DNA at A/T-rich sites. EMBO J. 8, 4189–95 (1989).

42. Khadake, J. R. & Rao, M. R. S. Condensation of DNA and Chromatin by an SPKK-Containing Octapeptide Repeat Motif Present in the C-Terminus of Histone H1 t Biochemistry 36, 1041–1051 (1997).

43. Papareddy, R. K. et al. Chromatin regulates expression of small RNAs to help maintain transposon methylome homeostasis in Arabidopsis. Genome Biol. 21, 251 (2020).

44. Sanulli, S. et al. HP1 reshapes nucleosome core to promote phase separation of heterochromatin. Nature 575, 390–394 (2019).

45. Ninova, M., Fejes Tóth, K. & Aravin, A. A. The control of gene expression and cell identity by H3K9 trimethylation. Development 146, dev181180 (2019).

46. Schmucker, A. et al. Crosstalk between H2A variant-specific modifications impacts vital cell functions. bioRxiv 2021.01.14.426637 (2021). doi:10.1101/2021.01.14.426637

47. Fauser, F., Schiml, S. & Puchta, H. Both CRISPR/Cas-based nucleases and nickases can be used efficiently for genome engineering in Arabidopsis thaliana. Plant J. 79, 348–359 (2014).

48. Clough, S. J. & Bent, A. F. Floral dip: a simplified method forAgrobacterium-mediated transformation ofArabidopsis thaliana. Plant J. 16, 735–743 (1998).

49. R Core Team. R: A language and environment for statistical computing. R Found. Stat. Comput. Vienna, Austria. URL http://www.R-project.org/. (2017).

50. Bernatavichute, Y. V, Zhang, X., Cokus, S., Pellegrini, M. & Jacobsen, S. E. Genome-wide association of histone H3 lysine nine methylation with CHG DNA methylation in Arabidopsis thaliana. PLoS One 3, e3156 (2008).

51. Krueger, F. Trim Galore. http://www.bioinformatics.babraham.ac.uk/projects/ (2019).

52. Dobin, A. et al. STAR: ultrafast universal RNA-seq aligner. Bioinformatics 29, 15–21 (2013).

53. Broad Institute. Picard Toolkit. Broad Institute, GitHub repository http://broadinstitute.github.io/picard/ (2019).

54. Anders, S., Pyl, P. T. & Huber, W. HTSeq--a Python framework to work with high-throughput sequencing data. Bioinformatics 31, 166–169 (2015).

55. Cheng, C.-Y. et al. Araport11: a complete reannotation of the Arabidopsis thaliana reference genome. Plant J. 89, 789–804 (2017).

56. Love, M. I., Huber, W. & Anders, S. Moderated estimation of fold change and dispersion for RNA-seq data with DESeq2. Genome Biol. 15, 550 (2014).

57. Lu, Z., Hofmeister, B. T., Vollmers, C., DuBois, R. M. & Schmitz, R. J. Combining ATAC-seq with nuclei sorting for discovery of cis-regulatory regions in plant genomes. Nucleic Acids Res. 45, e41–e41 (2017).

58. Buenrostro, J. D., Wu, B., Chang, H. Y. & Greenleaf, W. J. ATAC-seq: A Method for Assaying Chromatin Accessibility Genome-Wide. Curr. Protoc. Mol. Biol. 109, (2015).

59. Langmead, B. & Salzberg, S. L. Fast gapped-read alignment with Bowtie 2. Nat. Methods 9, 357–359 (2012).

60. Gaspar, J. M. Genrich. https://github.com/jsh58/Genrich (2019).

61. Ramírez, F., Dündar, F., Diehl, S., Grüning, B. A. & Manke, T. deepTools: a flexible platform for exploring deep-sequencing data. Nucleic Acids Res. 42, W187–W191 (2014).

62. Yelagandula, R., Osakabe, A., Axelsson, E., Berger, F. & Kawashima, T. Genome-Wide Profiling of Histone Modifications and Histone Variants in Arabidopsis thaliana and Marchantia polymorpha. in Methods in Molecular Biology 93–106 (2017). doi:10.1007/978-1-4939-7003-2_7

63. Hale, C. J. et al. Identification of Multiple Proteins Coupling Transcriptional Gene Silencing to Genome Stability in Arabidopsis thaliana. PLoS Genet. 12, 1–20 (2016).

64. Potok, M. E. et al. Arabidopsis SWR1-associated protein methyl-CpG-binding domain 9 is required for histone H2A.Z deposition. Nat. Commun. 10, 3352 (2019).

